# The Optimum Cut-off Value of the Nelwan Score to Diagnose Typhoid Fever in Adults

**DOI:** 10.1101/2021.02.22.432239

**Authors:** Erni Juwita Nelwan, Luh Putu Listya Paramita, Robert Sinto, Fransiscus Nikodemus Hosea, Pringgodigdo Nugroho, Herdiman T. Pohan

## Abstract

**Introduction:** Typhoid fever can be challenging to diagnose since clinicians often depend merely on clinical presentation. Clinical scores are useful to provide more accurate diagnosis. Variables in Nelwan Score are derived from clinical signs and symptoms of suspected cases for typhoid. Diagnostic value of Nelwan Score based on a cut-off value has never been evaluated.

**Methods:** A cross sectional study was conducted between July 2017 and January 2018 in five hospitals and two Primary Health Centers in Jakarta and Tangerang. The inclusion criteria were patients with 3-14 days of fever and gastrointestinal symptoms between July 2017 and January 2018. Diagnosis are confirmed by blood culture, rectal swab culture, or PCR. Cut-off analysis was performed by using Receiver Operating Characteristic (ROC) curve and diagnostic value was analyzed to generate sensitivity, specificity, predictive value and likelihood ratio.

**Result:** From 233 subjects involved, 4.72% of them were confirmed to have typhoid fever. The optimal cut-off value of Nelwan Score is 10 with AUC 71.3%. This cut-off value has sensitivity 81.8%, specificity 60.8%, PPV 9.3%, NPV 98.5%, LR + 2.086, and LR – 0.299.

**Conclusion:** Nelwan Score with cut-off value of 10 provides a good diagnostic performance as a screening tool for patients with suspected typhoid fever clinical presentation.

## INTRODUCTION

Typhoid fever caused by *Salmonella typhi* and *Salmonella paratyphi,* is a communicable disease inflicting a heavy public health burden. It is estimated that typhoid fever is responsible for over 22 million cases globally with more than 200,000 deaths in the year 2000, with the majority of cases reported from South and South-East Asia.^1,2^ In Indonesia, typhoid fever is endemic and needs serious attention because of the increasing incidence annually (500/100.000 population) and mortality rate (0.6-5%). The typhoid carrier screening among Jakarta’s food vendors in 2013 showed that 2.9% of them were typhoid carrier.^3^

The clinical presentation of typhoid fever adds to diagnostic difficulties since it mimics other febrile illnesses like malaria, dengue fever and influenza^4,5,6,7^ Nonetheless, the characteristics presentation features such as the *step ladder fever,* relative bradycardia, and coated tongue is considered to aid clinical diagnosis. These features were described by Haq et al to have provided a good specificity (100%, 94% and 94% respectively).^4^

The laboratory investigations (such as serological tests, culture, and PCR) which are used for screening and diagnosis, despite its role, still have substantial limitation on reliability, sensitivity and specificity. Additionally, laboratory investigations are not always available nor affordable, and sometimes contribute to delayed diagnosis and inappropriate treatment.^3,8,9,10^.

Despite, lots of effort being exercised to support different ways of diagnosis and management of typhoid fever, the incidence, morbidity, and mortality rates remain high.^2,3,11,12^ Therefore an early accurate diagnosis will help to decrease the typhoid fever burden including a decrease in laboratory investigations, antimicrobial resistance, and cost.

A clinical scoring system to aid the diagnosis of typhoid fever was first developed by Nelwan RHH in 1991 and is known as the Nelwan score.^13^ Accordingly, score points are given to the signs and symptoms, obtained from history and physical examination.The points obtained are totaled and the score corresponds to the probability of clinical diagnosis of typhoid fever: a score of 13 or more is rated highly likely to be positive and a score below seven to be less indicative of typhoid fever being positive. While the Nelwan Score can benefit the diagnosis of typhoid fever, the diagnostic value of it has not been evaluated, particularly compared with blood culture and PCR as the reference standard of typhoid diagnosis. This study aimed to determine the cut-off value of the Nelwan score in diagnosing typhoid fever in adult patients.

## METHODS

A cross-sectional study was conducted between July 2017 to January 2018 using primary data of patients who visited the Emergency department, Outpatient clinic and Internal Medicine ward in five hospitals (Persahabatan Hospital, Budhi Asih Hospital, South Tangerang Hospital, Hermina Ciputat Hospital, MMC Hospital) and two Primary Health Centers (Jatinegara and Gambir Primary Health Center) in Jakarta area. The inclusion criteria were adult patients (age from 18-65 years old) with fever ≥ 37,5°C ranging from 3 to 14 days and at least one abdominal manifestation indicative of typhoid fever. The exclusion criteria were pregnant women, those with known causes of fever, and those treated with antibiotics.

Clinical history and physical examination was performed on all study participants together with laboratory investigations (complete blood count, differential count, blood culture, PCR, and rectal swab). A written informed consent was obtained from all participants prior to data collection. Ethical approval was obtained from the Ethical Board of the Universitas Indonesia (Approval Number: 641/UN2.F1/ETIK/2017). All data collected were kept confidential.

Clinical manifestations, such as fever and abdominal symptoms were used to diagnose typhoid fever, and the clinical diagnosis was confirmed by positive blood culture or rectal swab culture or PCR. Descriptive analysis was performed on demographic data and clinical manifestation. Cut-off analysis was performed by using ROC curve with 95% confidence intervals (CI) and diagnostic value of Nelwan Score was analysed to generate sensitivity, specificity, predictive value and likelihood ratio.

### Nelwan Score

The score is calculated by giving a point to each of the following symptoms: fever less than one week, headache, weakness, nausea, abdominal pain, anorexia, vomiting, disturbed gastrointestinal motility, insomnia, hepatomegaly, and splenomegaly (Table 1). A two-score point is given for fever more than one-week, relative bradycardia, typhoid tongue, melena stools, and altered mental status, respectively. The maximum score obtained can be twenty. A score of 13 or more indicates that a patient is highly likely to be positive for typhoid fever, while a score ranging between 8 to 12 provides a 50% probability of typhoid fever positivity, and a score of 7 or less is less indicative of typhoid fever.

**Table 1.** Nelwan score showing the score allocation

| No | Sign and/or symptom | Score |
| --- | --- | --- |
| 1 | Fever < 1 week | 1 |
| 2 | Headache | 1 |
| 3 | Weakness | 1 |
| 4 | Nausea | 1 |
| 5 | Abdominal pain | 1 |
| 6 | Anorexia | 1 |
| 7 | Vomiting | 1 |
| 8 | Disturbed GI motility | 1 |
| 9 | Insomnia | 1 |
| 10 | Hepatomegaly | 1 |
| 11 | Splenomegaly | 1 |
| 12 | Fever >1 week | 2 |
| 13 | Relative bradycardia | 2 |
| 14 | Typhoid tongue | 2 |
| 15 | Melena stools | 2 |
| 16 | Impaired consciousness | 2 |
|  | Total score | 20 |

## RESULT

A total of 233 participants were enrolled and examined using both Nelwan score and the reference standard. The signs and symptoms reported by the participants are shown in Table 2. Proportion of typhoid cases with laboratory confirmation either by positive culture or positive PCR was 11/233 (4.72%). There were 123 (52.8%) women in total and 67 (54.5%) women among laboratory confirmed cases The proportion of women from the typhoid-fever-confirmed group were 54.5%. The median age of the study participants was 38 years, while the median age of typhoid-fever-confirmed cases was 26 years.

**Table 2.** Characteristics of study subjects (clinical and confirmed cases)

| Clinical characteristics | N = 233 | Confirmed cases | Clinical cases |
| --- | --- | --- | --- |
|  | n(%) | n | n |
| Gender |  |  |  |
| Male | 110 (47.2) | 50 | 60 |
| Female | 123 (52.8) | 67 | 56 |
| Age |  | 26 (20-45)* | 38 (18-81)* |
| Fever ≤ 1 week | 202 (86.7) | 11 (100) | 191 (86) |
| Headache | 209 (89.7) | 11 (100) | 198 (89.2) |
| Malaise | 221 (94.8) | 11 (100) | 210 (94.6) |
| Nausea | 222 (95.3) | 11 (100) | 211 (95) |
| Abdominal pain | 207 (88.8) | 11 (100) | 196 (88.3) |
| Anorexia | 211 (90.6) | 11 (100) | 200 (90.1) |
| Vomiting | 168 (72.1) | 10 (90,9) | 160 (72.1) |
| Motility disorder | 177 (76) | 11 (100) | 166 (74.8) |
| Insomnia | 174 (74.7) | 11 (100) | 163 (73.4) |
| Hepatomegaly | 7 (3) | 0 | 7 (3.2) |
| Splenomegaly | 0 | 0 | 0 |
| Relative bradycardia | 21 (9) | 7 (63.6) | 14 (6.3) |
| Typhoid tongue | 94 (40.3) | 10 (90.9) | 84 (37.8) |
| Melena | 21 (9) | 0 | 21 (9.5) |
| Decrease of consciousness | 7 (3) | 1 (9,1) | 6 (2.7) |
| Nelwan Score | 9 (4-15)* | 11 (9-13)* | 9 (4-15)* |

\*Median (min-maks)

The symptoms of fever for less than a week, headache, weakness, nausea, abdominal pain, anorexia, motility disorder, and insomnia were present (100%) in all the laboratory confirmed cases.

The figure 1 shows the ROC curve analysis and the diagnostic prediction according to the area under the curve (AUC) was 77.3% (95% CI 65.9% – 88.7%). The corresponding cut-off value to estimate the typhoid fever diagnosis and risk in patients with fever is shown in Figure 2. The cut off score was 8 considering there were no subjects in the fever group whose score was under 9.

**Figure 1.**
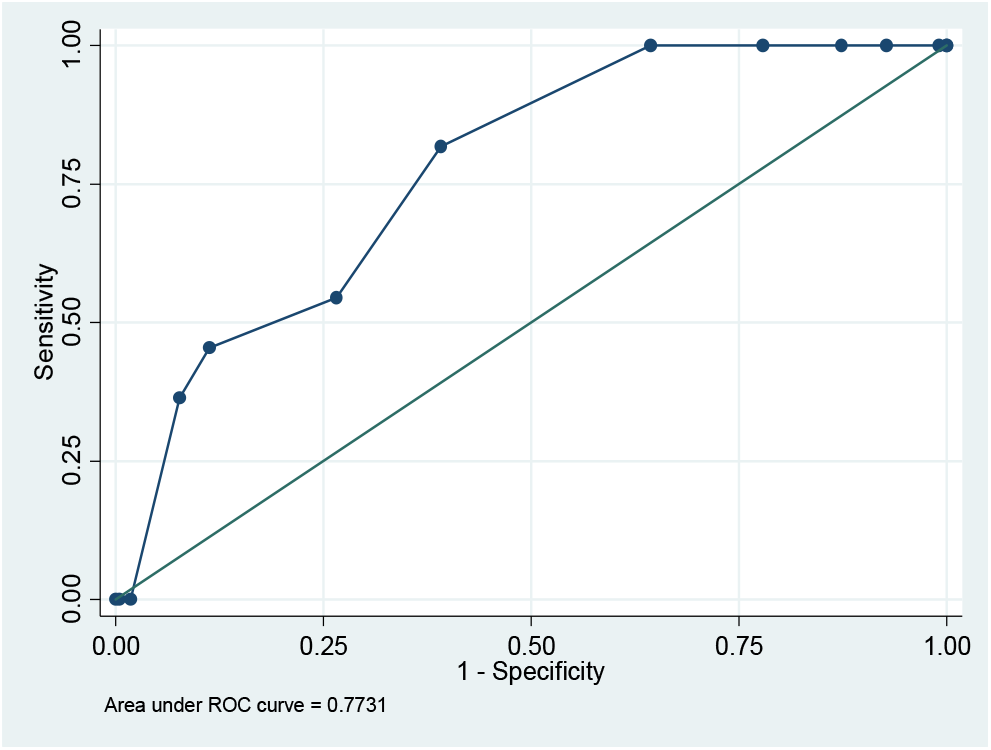
ROC (*Receiving Operator Characteristics*) Curve of Nelwan Score

**Figure 2.**
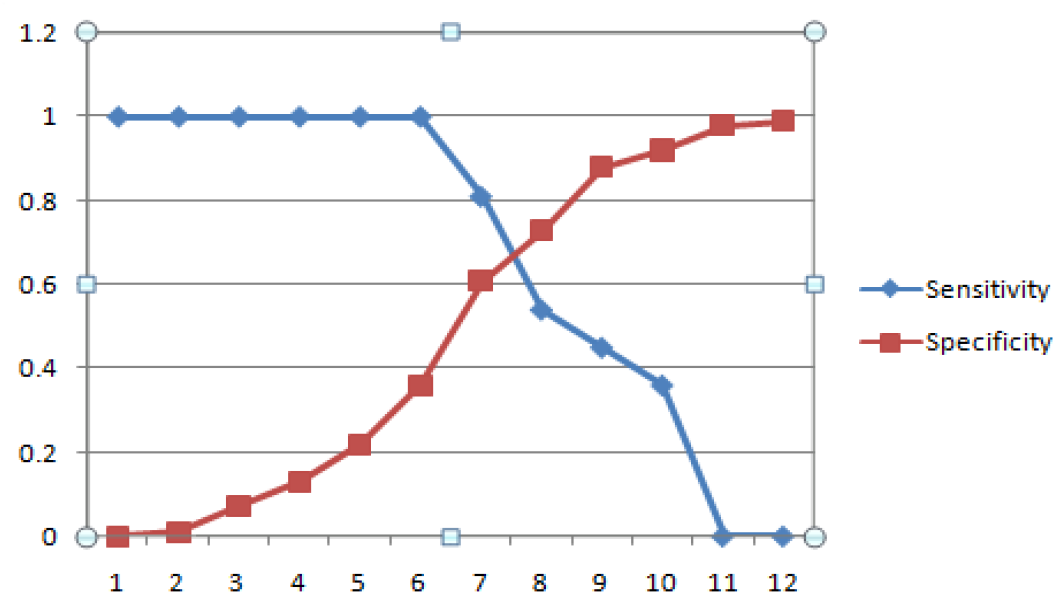
The curve showing the intersection between sensitivity and specificity of Nelwan Score

The Nelwan score was further explored with different cut-off values (Table 3). The cut-off value of 10 had the highest sensitivity (81.8*%*) with negative predictive value of 98.5% and reliability of 42.6%, while specificity of 88.7% was observed at cut-off value of 12.

**Table 3.** Diagnostic Value in Several Cut-off Points in Nelwan Score

| Cut- off Value | Sensitivity (%) (95%CI) | Specificity (%) (95%CI) | PPV(%) (95%CI) | NPV(%) (95%CI) | LR+ (95%CI) | LR- (95%CI) | Reliability (%) |
| --- | --- | --- | --- | --- | --- | --- | --- |
| 10 | 81.8<br>(52 - 95) | 60.8<br>(54-67) | 9.3<br>(5-17) | 98.5<br>(95- 100) | 2.086<br>(1.66-2.62) | 0.299<br>(0.19-0.47) | 42.6 |
| 11 | 54.5<br>(28-79) | 73.4<br>(67- 79) | 9.2<br>(4-19) | 97<br>(93-99) | 2.05<br>(1.47-2.87) | 0.62<br>(0.47-0.82) | 27.9 |
| 12 | 45.5<br>(21-72) | 88.7<br>(84-92) | 16.7<br>(7-34) | 97<br>(94-99) | 4.04<br>(2.33-7) | 0.61<br>(0.44-0.86) | 34.2 |

PPV : Positive Predictive Value, NPV : Negative Predictive Value, LR :Likelihood Ratio

## DISCUSSION

The usefulness of the Nelwan score for the diagnosis of typhoid fever was evaluated based on the clinical signs and symptoms of patients who presented to five hospitals and two community health facilities in greater Jakarta. We found the score of 10 corresponded to an acceptable sensitivity (81.8%) and negative predictive value (98.5) and it can be considered the cut-off value of the Nelwan score for diagnosing typhoid fever. These values indicate that the Nelwan score has practical use as a screening tool which would benefit clinicians working in health facilities with limited laboratory facilities.^14,15^ A tool that aids early diagnosis and prompt treatment of typhoid fever is important in Indonesia to reduce the morbidity and prevention of new typhoid carriers. Particularly in dealing with typhoid fever, accurate early diagnosis and prompt treatment are important to reduce morbidity and prevent new carrier cases.

We had used the ROC curve to evaluate the Nelwan score, since AUC are used to determine the usefulness of the test. The value of AUC (77.3% (95% CI, 65.9% – 88.7%) is statistically considerable and gives the best trade-off between true positives and lowest false positives., Since there were no previous studies on Nelwan Score to assess cut-off value, we were unable to compare the discriminatory ability of our findings. Nonetheless, our findings suggest that a patient with clinical symptoms suspected of typhoid fever has a Nelwan score of ≥ 10, clinicians could initiate treatment promptly where investigations such as cultures are not available. In cases, where the Nelwan score is ≤ 10, laboratory investigations should be performed to confirm the diagnosis.

A further analysis was performed using different cut-off values (11 and 12) to explore the diagnostic accuracy of the Nelwan score. Despite the higher values, the sensitivity was lower for scores of 11, and 12 than for the score of 10, while specificity increased. In addition to sensitivity and specificity the possibility that a patient has typhoid fever at a score of 10 is indicated by the positive and negative likelihood ratio values. Our findings compare with the findings reported by Hargo et al, and Haq who suggested that clinical features were a strong marker in cases of typhoid fever. Likewise, several other studies have indicated the benefit of clinical features together with laboratory tests for the diagnosis of typhoid fever. ^4,5,6,16^ Another criteria for the diagnosis of enteric fever was developed by Neopane et al. It primarily differentiates the clinical manifestation into three major (headache, fever, and relative bradycardia) and five minor criteria (abdominal pain, vomiting, diarrhea, splenomegaly and chill) and is therefore different from the Nelwan scoring system. Although their specificity was higher, the sensitivity was lower than the value we found for our score of 10. Thus, Nelwan score will be useful to screen for typhoid fever where sophisticated laboratory tests such as blood/stool cultures are not available.

Statistically, patients with a total score of 10 could be estimated to have a confirmed typhoid fever case with sensitivity, specificity, positive predictive value, and negative predictive value of 81.8%, 60.8%, 9.3%, and 98.5%, respectively. Positive likelihood ratio and negative likelihood ratio of Nelwan Score in this setting were 2.086 and 0.299. Positive predictive value was very low because of the low proportion of typhoid fever case in this study. We also attempted to obtain a higher sensitivity value, where sensitivity was estimated to increase up to 90% or more. But, in cut-off points of 11 and 12, we found there was no significant increase of sensitivity, specificity, and AUC value.

When the cut-off value was 10, the AUC value was 71.3, which showed a good clinical discrimination. Based on its sensitivity and its high negative predictive value, Nelwan Score might hold a valuable role as a screening tool in patients with suspected typhoid fever. If a patient with suspected typhoid fever has Nelwan Score less than 10, the probability of the patient to suffer from typhoid fever is low. In this condition, the clinician using Nelwan Score should consider other etiologic agents. Meanwhile when a patient with suspected typhoid fever has Nelwan Score ≥ 10, further investigation needs to be performed to support the diagnosis.

Hargo et al, used secondary data and TUBEX TF as reference, reported that Nelwan Score had a sensitivity value of 94%, specificity of 87%, and accuracy of 90%.^16^ Supporting the functionality of Nelwan Score, which based on clinical manifestation, Haq et al reported that several clinical features serve as strong markers of typhoid fever. They found that stepladder pattern of rise in temperature, loose bowel movements, relative bradycardia and a coated tongue had a high specificity (>90%) for typhoid fever.^4^ Similarly, other studies by Kuvandik et al, Hosoglu et al, and Hadianti et al also showed that clinical manifestation and laboratory tests altogether can be beneficial in diagnosing enteric fever in the absence of confirmatory tests.^5,6,17^ Consistently, Ross et al showed that clinical features are useful in diagnosing enteric fever when culture is not available.^15^ Several studies mentioned before showed that clinical manifestation serves an important diagnostic value, but most of them also included laboratory workups as components of their diagnostic prediction model.

Nelwan Score which also focuses primarily on clinical manifestation, on the other hand, has a better sensitivity (81.8%). This makes Nelwan Score serves as one of the best available clinical screening tools in patients with suspected typhoid fever. Furthermore, this score can be extensively implemented in a lot of healthcare facilities with limited resource as a typhoid fever screening tool.

In our study, the proportion of confirmed typhoid fever cases was low (4.72%). Although this may not be a representative figure of typhoid fever in Indonesia, it is similar to the proportion reported by the study conducted in north Jakarta hospitals by Punjabi et al.^18^ The low incidence might have contributed to the lower positive predictive values in our study. Another limitation is that cultures both blood and rectal swab taken only at one time point may have limited diagnostic sensitivity.^19^ Similar factors apply to PCR.^20^ Meanwhile doing confirmatory investigation at different times in the course of the illness was not possible in our study setting. Still and all, this number had not yet represented the real proportion of typhoid fever in Indonesia. Punjabi et al, involving subjects with 3-day-or-more fever accompanied by abdominal complaints in North Jakarta hospitals, also found the similarly small proportion. The small proportion result in their study was due to passive case detection, which made typhoid fever patients who were not coming to the hospitals in their study area remained unknown.^18^ Additionally, Al Emran et.al also reported that only a small proportion of typhoid fever cases that can be detected by blood culture. In their study, antibiotics administration and duration between onset of disease and sample collection contributed to the small number of typhoid cases.^20^

Another reason why this study only reported a small proportion of typhoid fever case is that cultures (especially rectal swab culture) have a limited sensitivity which varies throughout the course of disease (40-80% during incubation period) and at the same time influenced by antibiotics administration.^18,19^ The same factors also apply to PCR. Although some studies showed that PCR has a high sensitivity and specificity, false negative may still occur.^20,21,22^ False negative in PCR may result from small numbers of bacteria, gene mutation, and inhibitory substances in the sample.^21,22,23,24^

## CONCLUSION

The clinical features used in the Nelwan Score showed the cut off value of 10 was the score corresponding to the best diagnostic accuracy values. Taking this into consideration, the Nelwan score is applicable as a screening tool for patients presenting with suspected typhoid fever features in limited resources settings.

## Acknowledgements

We acknowledge Dr. Rukhsana Ahmed for reviewing the final draft and thank the Division of Tropical and Infectious Disease Department of Internal Medicine Universitas Indonesia/Cipto Mangunkusumo Hospital, Persahabatan Hospital, Budhi Asih Hospital, South Tangerang Hospital, Hermina Ciputat Hospital, MMC Hospital, Jatinegara Primary Health Center and Gambir Primary Health Center for accommodating and supporting this research. We also thank the study subjects for willing participating in the study.

